# FZD10 regulates cell proliferation and mediates Wnt1 induced neurogenesis in the developing spinal cord

**DOI:** 10.1101/690263

**Authors:** Abdulmajeed Fahad Alrefaei, Andrea E. Münsterberg, Grant N. Wheeler

**Affiliations:** School of Biological Sciences, University of East Anglia, Norwich Research Park, Norwich, NR4 7TJ, UK; Jamoum University College, Department of Biology, University of Umm Al-Qura, Saudi Arabia

## Abstract

Wnt/FZD signalling activity is required for spinal cord development, including the dorsal-ventral patterning of the neural tube, where it affects proliferation and specification of neurons. Wnt ligands initiate canonical, β-catenin-dependent, signaling by binding to Frizzled receptors. However, in many developmental contexts the cognate FZD receptor for a particular Wnt ligand remains to be identified. Here, we characterized FZD10 expression in the dorsal neural tube where it overlaps with both Wnt1 and Wnt3a, as well as markers of dorsal progenitors and interneurons. We show FZD10 expression is sensitive to Wnt1, but not Wnt3a expression, and FZD10 plays a role in neural tube patterning. Knockdown approaches show that Wnt1 induced ventral expansion of dorsal neural markes, Pax6 and Pax7, requires FZD10. In contrast, Wnt3a induced dorsalization of the neural tube is not affected by FZD10 knockdown. Gain of function experiments show that FZD10 is not sufficient on its own to mediate Wnt1 activity *in vivo*. Indeed excess FZD10 inhibits the dorsalizing activity of Wnt1. However, addition of the Lrp6 co-receptor dramatically enhances the Wnt1/FZD10 mediated activation of dorsal markers. This suggests that the mechanism by which Wnt1 regulates proliferation and patterning in the neural tube requires both FZD10 and Lrp6.

## Introduction

Following neural tube formation from the neural plate, complex tissue interactions and signalling pathways contribute to its patterning and differentiation, to generate well defined neuronal populations along its dorsal ventral (DV) axis. The roof and floor plates are signaling centres that govern the formation of sensory neurons in the dorsal part and motor neurons in the ventral part. The roof plate secretes members of the Wnt and BMP families, whilst the floor plate produces Sonic hedgehog (Shh). These secreted signaling molecules are crucial for neural tube patterning along the dorso-ventral axis (reviewed in Le Dréau and Martí, 2012).

Wnt glycoproteins bind to Frizzled (FZD) receptors and Lrp5/6 co-receptors to initiate β-catenin/TCF-dependent activation of Wnt target genes in the nucleus (Nakamura et al., 2016; Moon et al., 2004; Logan and Nusse, 2004). Wnt proteins regulate cell proliferation and specification during nervous system development. In mice lacking Wnt1, the midbrain is lost and the hindbrain is affected. In Wnt3a knockout mice the anterior-posterior axis is truncated and the hippocampus is lost (reviewed in Amerongen and Berns, 2006). In Wnt1^−/−^ Wnt3a^−/−^ double mutant mice the specification of dorsal neurons is affected (Ikeya et al., 1997; Muroyama et al., 2002).

Several Wnt family members, including Wnt1 and Wnt3a, are expressed in the roof plate of the neural tube in chick and mouse, where they promote proliferation of neural progenitors (Hollyday et al., 1995; Chesnutt et al., 2004; Dickinson et al., 1994; Zechner et al., 2003; Ille et al., 2007; Bonner et al., 2008). Additionally, Wnt1 and Wnt3a are implicated in dorso-ventral (DV) patterning of the neural tube and co-overexpression of Wnt1 and Wnt3a in the chick neural tube results in activation of dorsal markers (Pax6/7) and repression of ventral markers (Olig2 and Nkx2.2) (Alvarez-Medina et al., 2008). It is still unclear, however, which FZD receptors mediate canonical Wnt1/3a signaling in the neural tube.

Genetic experiments have shown that FZD receptors are involved in neural tube development. For example, neural tube closure is affected in FZD1 and FZD2 knockout mice (Yu et al., 2010). The neural tube also fails to close in FZD3^−/−^ FZD6^−/−^ double mutants (Wang et al., 2006). FZD3 knockout mice show severe defects in axon development in the central nervous system and some neurons fail to migrate and cluster in the midline of the spinal cord (Hua et al., 2013; Wang et al., 2006). In addition the FZD co-receptor, Lrp6, which has been shown to bind Wnt1, is necessary for the activation of Wnt signaling (He et al., 2004; MacDonald and He, 2012; Tamai et al., 2000). Lrp6 is expressed in neural tube and its mutations result in neural tube defects including a failure of neural tube closure and disruption of cell polarity (Allache et al., 2014; Gray et al., 2013; Houston and Wylie, 2002). In chick embryos, expression of FZD receptors has been characterized during early development. FZD receptors are detected in different tissues, including the developing brain (Chapman et al., 2004; Fuhrmann et al., 2003; Stark et al., 2000; Quinlan et al., 2009; Theodosiou and Tabin, 2003).

FZD10 is one of the FZD family receptors that has been detected in different species, including zebrafish, Xenopus, chick and mouse (Galli et al., 2014; Kawakami et al., 2000a; Moriwaki et al., 2000; Wheeler and Hoppler, 1999; Nikaido et al., 2013; Yan et al., 2009). We previously showed that FZD10 is expressed in the dorsal neural tube (Wheeler and Hoppler, 1999) and using axis duplication assays in early *Xenopus* embryos, we showed that FZD10 acts through canonical Wnt signaling. In addition, a FZD10 knockdown phenotype was rescued by β-catenin injections, suggesting that β-catenin is downstream of FZD10 (Garcia-Morales et al., 2009).

Here we investigated the potential function of FZD10 as a mediator of canonical Wnt signaling in the developing chick neural tube. We examined FZD10 expression and its relationship with Wnt1 and Wnt3a in the dorsal neural tube. Using *in ovo* electroporations of shRNA, we show that FZD10 knockdown affects cell proliferation and differentiation of the neural tube. Targeted mis-expression of Wnt1 and Wnt3a show that Wnt1 positively affects FZD10 expression whereas Wnt3a has no effect, suggesting that Wnt1 may act through FZD10. Consistent with this idea, FZD10-shRNA inhibited the Wnt1-mediated dorsalization of the neural tube. To determine the importance of the Lrp6 co-receptor in Wnt1/FZD10 signaling *in vivo* we used co-electroporations into the neural tube. This revealed that Lrp6 enhances Wnt1/FZD10 mediated activation of dorsal markers during spinal cord neurogenesis. Luciferase reporter assays (TOP-flash) confirmed that FZD10 and Lrp6 are required for Wnt1 biological activity *in vivo*. This suggests the mechanism by which Wnt1 regulates proliferation and patterning of the developing spinal cord involves interactions with both FZD10 and Lrp6.

## Results

### FZD10 is expressed in the dorsal domain of the spinal cord during neurogenesis

To identify receptors that could potentially mediate Wnt signalling during spinal cord neurogenesis we performed expression analysis of multiple frizzled (FZD) receptors in chick embryos. Subsequent investigations focused on FZD10, and a detailed time course examined FZD10 expression before the onset of neurogenesis, during initiation of neurogenesis and during formation of dorsal neurons (Fig. 1). At HH12, before the onset of neurogenesis, FZD10 expression was graded from dorsal to ventral (Fig. 1A). FZD10 continued to be expressed in the spinal cord but expression became dorsally restricted during the initiation of neurogenesis (Fig. 1A, HH14-20). During neurogenesis (HH18-24), FZD10 was expressed in regions of the spinal cord where dorsal progenitors arise (Fig. 1A, HH18-24). High levels of FZD10 transcripts were seen in the ventricular zone where progenitors are still proliferating.

**Figure 1:**
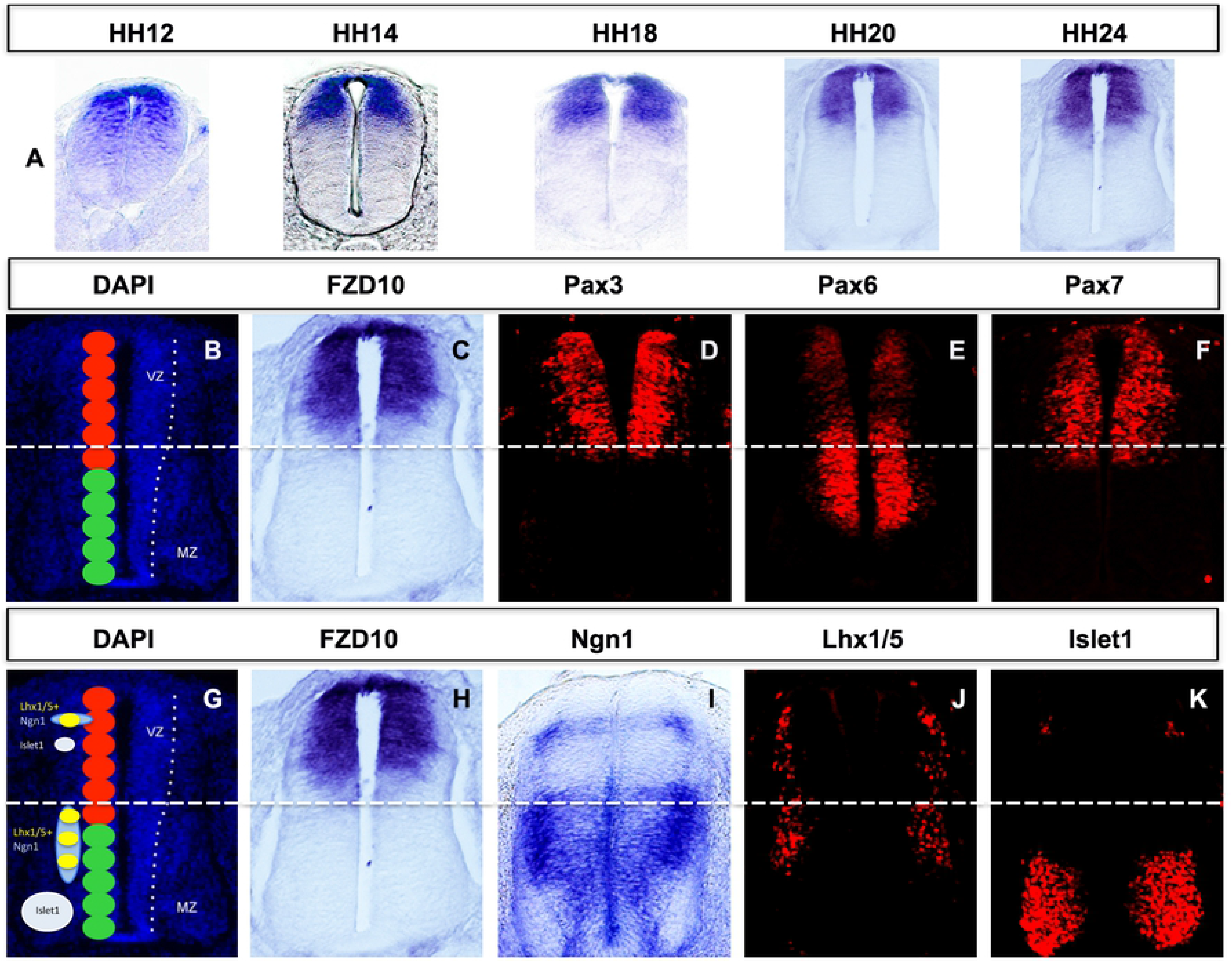
FZD10 expression in the neural tube correlates with markers of neural progenitors and differentiated neurons. (A) In situ hybridization shows the dorso-ventral extent of FZD10 expression from HH12-24. (B) Transverse section of the developing spinal cord at stage HH24 stained with DAPI; red circles represent 6 dorsal progenitor domains (dp 1-6) and green circles represent 5 ventral progenitor domains. (C) FZD10 transcript distribution compared with progenitor markers (D) Pax3, (E) Pax6 and (F) Pax7 detected by immunostain. (G) Schematic representation of differentiated neuron marker expression at HH24. (H) FZD10 expression compared with differentiated neurons markers detected by in situ hybridization, (I) Ngn1, or by immunostain, (J) Lhx1/5 and (K) Islet1. The same sections are shown in (C) and (H).

Based on the expression of specific transcription factors the dorsal ventral axis of the spinal cord can be divided into eleven domains. For example, Pax3/7 expression marks the dorsal progenitor domains dp1-6, and Pax6 marks dorsal progenitor domains, low expression in dp1-3 and higher expression in dp4-6, and one intermediate ventral progenitor domain, p0 (Le Dreau and Marti, 2012). Expression was compared to these well-characterized markers at stage HH24 (Fig. 1B-F and H-K), and FZD10 expression overlapped with Pax3 and Pax7 in progenitor domains dp 1-5 (Fig. 1B, C, D, F). In addition, FZD10 and Pax7 were expressed in the roof plate but Pax3 was not (Fig. 1C, D, F). In dorsal progenitor domains dp1-5, FZD10 overlapped with Pax6 expression which was weakly expressed there (Fig. 1B, C, E). FZD10 expression was seen in dorsal regions in which neural differentiation markers were expressed: Ngn1 (dp 2), Islet1 (dp 3) (Fig. 1G, H, I, K) and Lhx1/5 (dp 2-4) (Fig. 1J and K). In summary, FZD10 was strongly expressed in dorsal domains of the spinal cord that were positive for dorsal progenitor interneuron markers.

### FZD10 knockdown affects cell proliferation, dorso-ventral patterning and neurogenesis

To determine the requirement of FZD10 in spinal cord development three plasmids were commercially designed producing short-hairpin RNAs (shRNA) specifically against chick FZD10. FZD10 shRNA plasmids (pRFP-C-RS) were electroporated into one side of chick neural tubes at stage HH11-12. After 48 hours, embryos were screened for RFP expression and processed for phenotypic analysis by in situ hybridization. First, shRNA vectors were electroporated individually to assess FZD10 knockdown. Electroporation of FZD10 shRNA vectors B and C resulted in an overall reduction of endogenous FZD10 transcripts on the electroporated side of the spinal cord compared to the non-electroporated side. Although electroporation is mosaic and there is residual expression. Scrambled shRNA plasmids had no effect on expression of FZD10 (Supp Fig. 1).

Next, we analysed the effects of FZD10 knockdown on spinal cord development. Cryosections of embryos electroporated with FZD10 shRNA plasmids showed that the electroporated side was thinner with a shortened dorso-ventral axis (Fig. 2 D, E), suggesting that proliferation could be affected in the ventricular zone where neural progenitors are located. To confirm this, we used immunostaining for phospho-histone H3 (pH3). Quantification of the number of pH3 positive cells showed that the number of mitotic cells was reduced on the experimental side of the spinal cord after FZD10 knockdown (1.4 fold, p=0.01) (Fig. 2 F). Scrambled shRNA plasmids did not affect number of pH3 positive cells (Fig. 2 C). This showed that FZD10 knockdown by shRNA results in a reduction in cell polifration in the neural tube. Consistent with these results FZD10 knockdown by morpholinos reduced poliferation in dorsal regions of the neural tube in a previous report (Galli et al., 2014).

**Figure 2:**
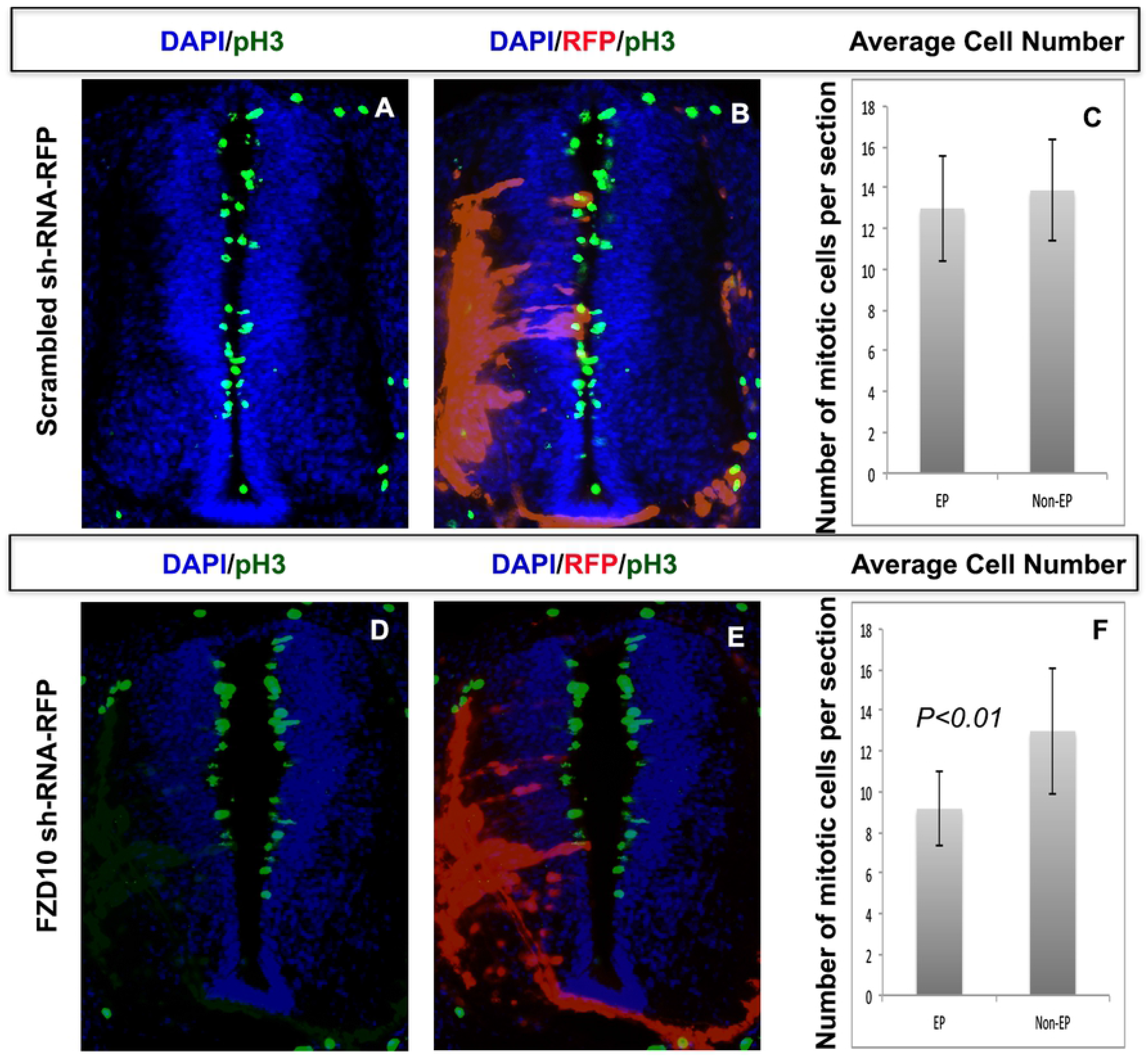
Knockdown of FZD10 results in a decrease in cell proliferation in the neural tube. (A, B, D, E) Cross sections of neural tube stained with DAPI and phosphor-histone H3 (green), showing the electroporated (RFP+ve) and non-electroporated sides. Electroporation with scrambled or FZD10 shRNA vectors as indicated. (C) Quantification of pH3 positive cells revealed that scrambled shRNA induced no significant difference in the number of proliferative cells on electroporated (EP) compared to the non-electroporated (Un-EP) side of the neural tube. (F) Quantification of the number of pH3 positive cells per section in the neural tube electroporated with FZD10 shRNA vector. Compared to the non-electroporated side there was a statistically significant decrease in the number of pH3 positive cells (students t-test (paired).

To examine effects of FZD10 knockdown on dorsal-ventral patterning and neural differentiation, we used markers that overlap with regions of FZD10 expression. FZD10 shRNA vectors were electroporated into one side of the neural tube at HH11-12, followed by incubation for 24 or 48 hours. Cryosections were immunostained for RFP, Pax6 and Pax7. After 24 hours post-electroporation the Pax7 expression domain was shifted dorsally on the FZD10 shRNA transfected side compared to the control side (Supp Fig. 2D-F). Similar observations were made after 48 hours post-electroporation of FZD10-shRNA; the expression domains of Pax6 and Pax7 were dorsally restricted on the electroporated side compared to the control side (12/13) (Fig. 3B, D). Scrambled shRNA electroporation had no effect on expression of Pax7 or of other markers (Supp Fig. 2 A-C, Fig. 3 A, C). Next, we assessed effects of FZD10 knockdown on spinal cord neurogenesis. After electroporation with scrambled or FZD10 shRNA vector into neural tubes at stage HH11-12, embryos were immunostained for differentiated neuron markers Lhx1/5 and Tuj-1. After 48 hours, Lhx1/5 expression was strongly reduced on the electroporated side of the spinal cord compared to the control (n=7/8) (Fig. 3F-F”), scrambled shRNA plasmid had no effect (Fig. 3E, E’). Tuj-1 expression was reduced 24 hours after electroporation with FZD10 shRNA on the electroporated side (Supp Fig. 2H). A reduction of Tuj-1 expression was also evident 48 hours post-electroporation, especially in the dorsal domain when compared with the control side (7/8) (Fig. 3H-H’’). Area measurements using ImageJ/Fiji showed that areas of expression were reduced for Pax7, Pax6, lhx1/5 and Tuj1 on the FZD10-shRNA electroporated side (Fig. 3B”, D”, F”, H”). Thus electroporation of FZD10 shRNA vectors inhibited cell proliferation in the ventricular zone and therefore affected dorso-ventral spinal cord patterning and neurogenesis.

**Figure 3:**
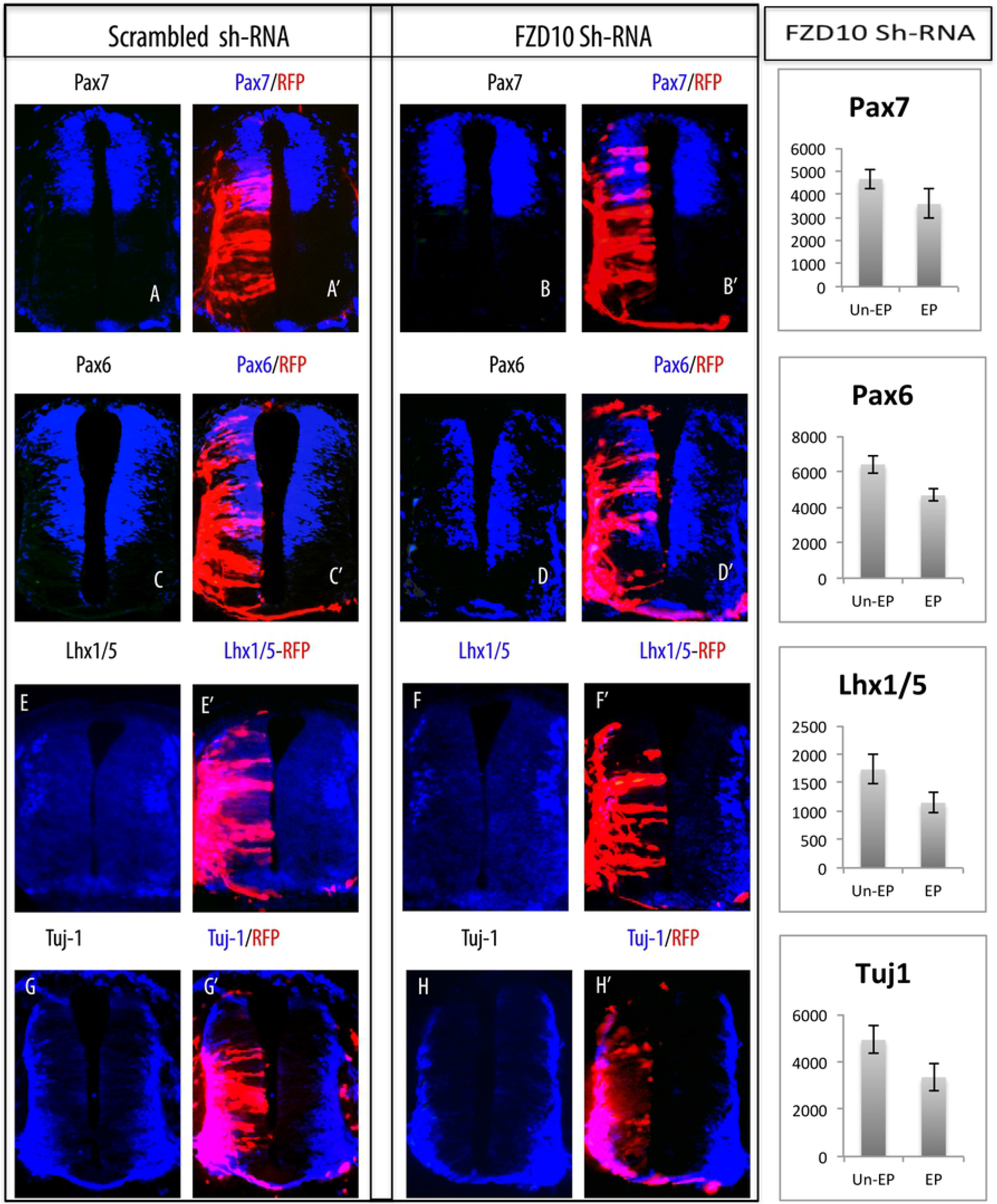
Effects of FZD10 knockdown on neurogenesis 48hours post-electroporation. Scrambled or FZD10 shRNA vectors were electroporated into neural tubes, these vectors expressed RFP to indicate successful transfection. (A,A’) Pax7 expression was identical on both sides after scrambled shRNA-vector electroporation. (B,B’) The ventral extend of the Pax7 domain was reduced on the electroporated side after FZD10 shRNA transfection. (C,C’) Pax6 expression was not affected after scrambled shRNA. (D, D’) But Pax6 expression was shifted dorsally on the electroporated side of the spinal cord after electroporation with the FZD10 shRNA-vector. (E, E’) Lhx1/5 expression in embryos electroporated with scrambled shRNA. (F, F’) Lhx1/5 expression was reduced after FZD shRNA-vector electroporation. (G, G’) Tuj-1 expression after scrambled shRNA electroporation. (H,H’) Tuj-1 expression was reduced in the spinal cord electroporated with FZD10 shRNA-vector. (B’’, D’’, F’’ and H’’) ImageJ was used to measure and compare the areas of expression on both sides of the spinal cord after FZD10 shRNA electroporation; neural marker expression was reduced on electroporated sides (EP).

### Wnt1 regulates FZD10 expression in the developing spinal cord

Wnt1 and Wnt3a are known to be involved in proliferation, neural specification and dorsal-ventral patterning of chick and mouse neural tube, and targeted misexpression of Wnt1 and Wnt3a leads to a ventral expansion of dorsal markers (Alvarez-Medina et al., 2008; Megason et al., 2002; Muroyama et al., 2002). First, we recapitulated these results (Supp Figs. 3, 4). Next, to investigate whether this is mediated by FZD10, we determined that FZD10 was co-expressed with Wnt1 and Wnt3a in the dorsal neural tube and roof plate (Supp Fig. 5). At stage HH14, FZD10 expression overlapped with Wnt1 and Wnt3a in the dorsal domain of the neural tube. By HH20, Wnt1 and Wnt3a expression was dorsally restricted whilst FZD10 expression still extended across the dorsal part of the spinal cord, consistent with a previous report (Galli et al., 2014). We next asked if FZD10 expression is affected by Wnt1 and Wnt3a electroporation in the neural tube. Embryos were electroporated at HH11-12 and after 48hours GFP indicated the transfected area. In situ detection of FZD10 transcripts after Wnt1 transfection revealed its broader and ventrally extended expression with strong signal in the roof plate (n=13/15) (Fig. 4A, B). However, FZD10 expression was not affected by Wnt3a (n=12/15) (Fig. 4C, D), suggesting that Wnt1 but not Wnt3a regulates expression of FZD10 in the dorsal neural tube.

**Figure 4:**
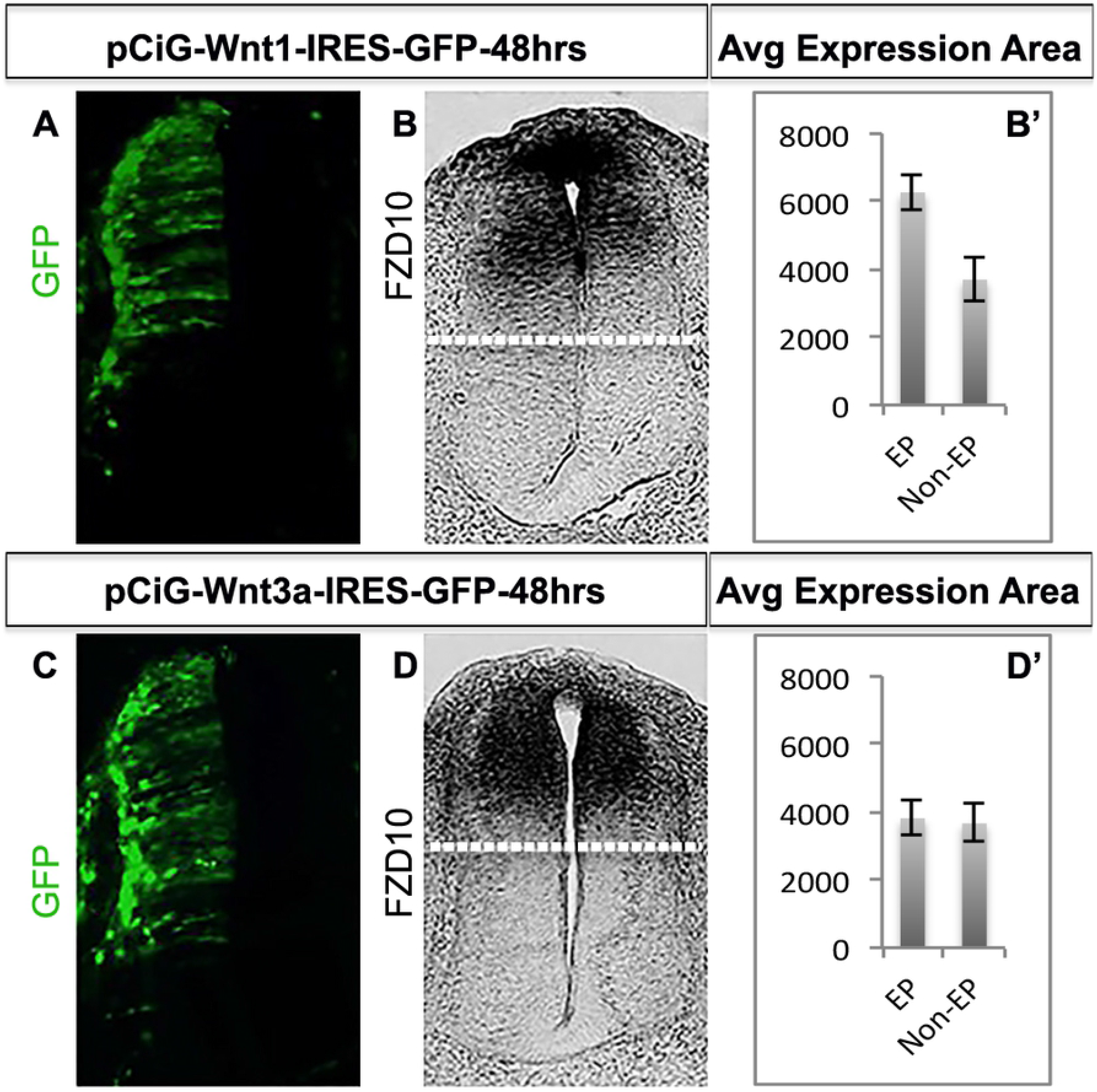
FZD10 expression expands ventrally following Wn1 misexpression. (A) GFP expression on the electroporated side indicating ectopic Wnt1 expression. (B) FZD10 expression was ventrally expanded on the transfected side as shown by (B’) area measurements. (C) GFP expression indicates ectopic Wnt3a expression. (D) FZD10 expression was unchanged on the transfected side as shown by (D’) area measurements.

### FZD10 is required for dorsalization of the neural tube in response to Wnt1 but not Wnt3a

Previous work showed that FZD10 mimics canonical Wnt activity and results in axis duplication in early *Xenopus* embryos (Garcia-Morales et al., 2009). Since Wnt1 promoted expression of FZD10, we wondered whether FZD10 is required for Wnt1 dependent dorsal patterning of the neural tube. To address this, we assessed whether shRNA-mediated FZD10 knockdown could rescue the effects of Wnt1 or Wnt3a overexpression on dorsal neural tube patterning. Embryos were co-electroporated at HH11/12 with Wnt1 or Wnt3a expression vectors and scrambled or FZD10 shRNA vectors (Fig 5). Co-electroporation of scrambled shRNA vectors with Wnt1 (n=8/8) or Wnt3a (n=6/6) had no effect on the ventral expansion of neural markers Pax6 and Pax7 after 48 hours (Fig. 5 A, F, C, H, Supp Figs. 3, 4). In contrast, co-electroporation of FZD10 shRNA lessened the effect of Wnt1 overexpression (n=14/14) and expression domains of Pax6 and Pax7 on the electroporated side were comparable to the control side (Fig. 5 B, D). Interestingly, the Wnt3a-induced ventral expansion of Pax6/Pax7 was not affected by FZD10 knockdown (n=13/14) (Fig. 5 G, I). The effects of scrambled and FZD10 shRNA transfection on Wnt1 and Wnt3 mediated neural tube patterning were quantified by area measurements using ImageJ/Fiji (Fig. 5 E, J). This showed that FZD10 is required for Wnt1-dependent ventral expansion of dorsal neural tube markers, although a direct interaction between Wnt1 and FZD10 remains to be confirmed. The results also indicated that Wnt3a presumably acts through different FZD receptors.

**Figure 5:**
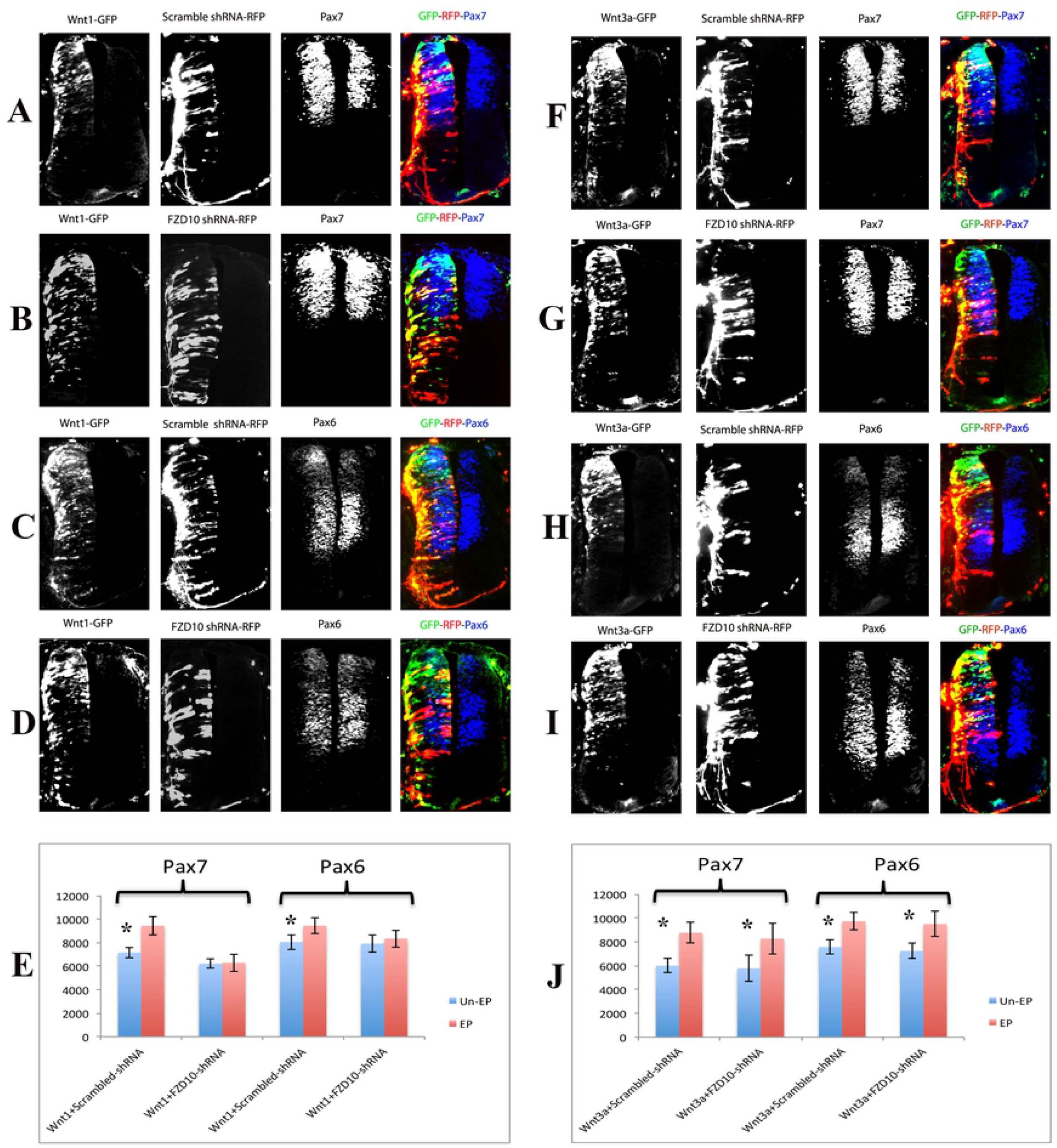
FZD10 mediates Wnt1-induced ventral expansion of dorsal neural tube markers. (A, C, E) Immunostaining showed that Pax6 and Pax7 expression was ventrally expanded after Wnt1 co-electroporation with scrambled shRNA. (B, D, E) Co-electroporation of FZD10 shRNA with Wnt1 inhibited the Wnt1-induced phenotype and abrogated the ventral expansion of Pax6 and Pax7 expression domains on the electroporated side. Co-electroporation of Wnt3a with (F, H) scrambled shRNA or (G, I) FZD10 shRNA had no effect on the Wnt3a induced phenotype and ventral expansion of Pax6 or Pax7 was evident on the electroporated sides of the spinal cord (J).

### Lrp6 co-receptor enhances FZD10 function during Wnt1 induced spinal cord dorsalization

Interestingly, FZD10 overexpression on its own did not lead to a ventral expansion of dorsal neural tube markers. Indeed, the expression of Pax6 and Pax7 was shifted dorsally and the ventral marker Nkx2.2 was expanded after electroporation of a FZD10 expression vector (Supp Fig. 6). This suggests FZD10 alone is not sufficient to dorsalize the neural tube. To explain the dorsal shift of Pax6 and Pax7 induced by FZD10 we tested whether receptor overexpression may restrict Wnt1 ligand, possibly by acting as a sponge. Consistent with this idea, FZD10 electroporation together with Wnt1 abrogated the Wnt1-induced phenotype and the ventral expansion of Pax7 was less pronounced compared to that seen after Wnt1 electroporation alone (Fig. 6 G, compare with Fig. 5 A). In addition, lengthening of the dorso-ventral axis was no longer evident; axis expansion often leads to a kink in the ventral part of the spinal cord and can be seen after overexpression of Wnt1 or Wnt3a (Fig. 6 E-H compare with Fig. 5 A, C). Furthermore, co-electroporation of FZD10 with LRP6, the frizzled co-receptor, or overexpression of LRP6 on its own, did not result in a ventral expansion of the dorsal marker Pax7 (n=8/10)(Fig. 6 A-D and data not shown). To test whether this could be due to a limited availability of endogenous Wnt1 ligand in the tissue we co-transfected FZD10, LRP6 and Wnt1 into the neural tube. This led to a dramatic lengthening of the axis and to ventral expansion of Pax7 expression compared to the control side (Fig. 6 I-L). Transfection of receptor and co-receptor, FZD10 and LRP6, together with Wnt1 enhanced the phenotype compared to Wnt1 alone (Supp Fig. 7).

**Figure 6:**
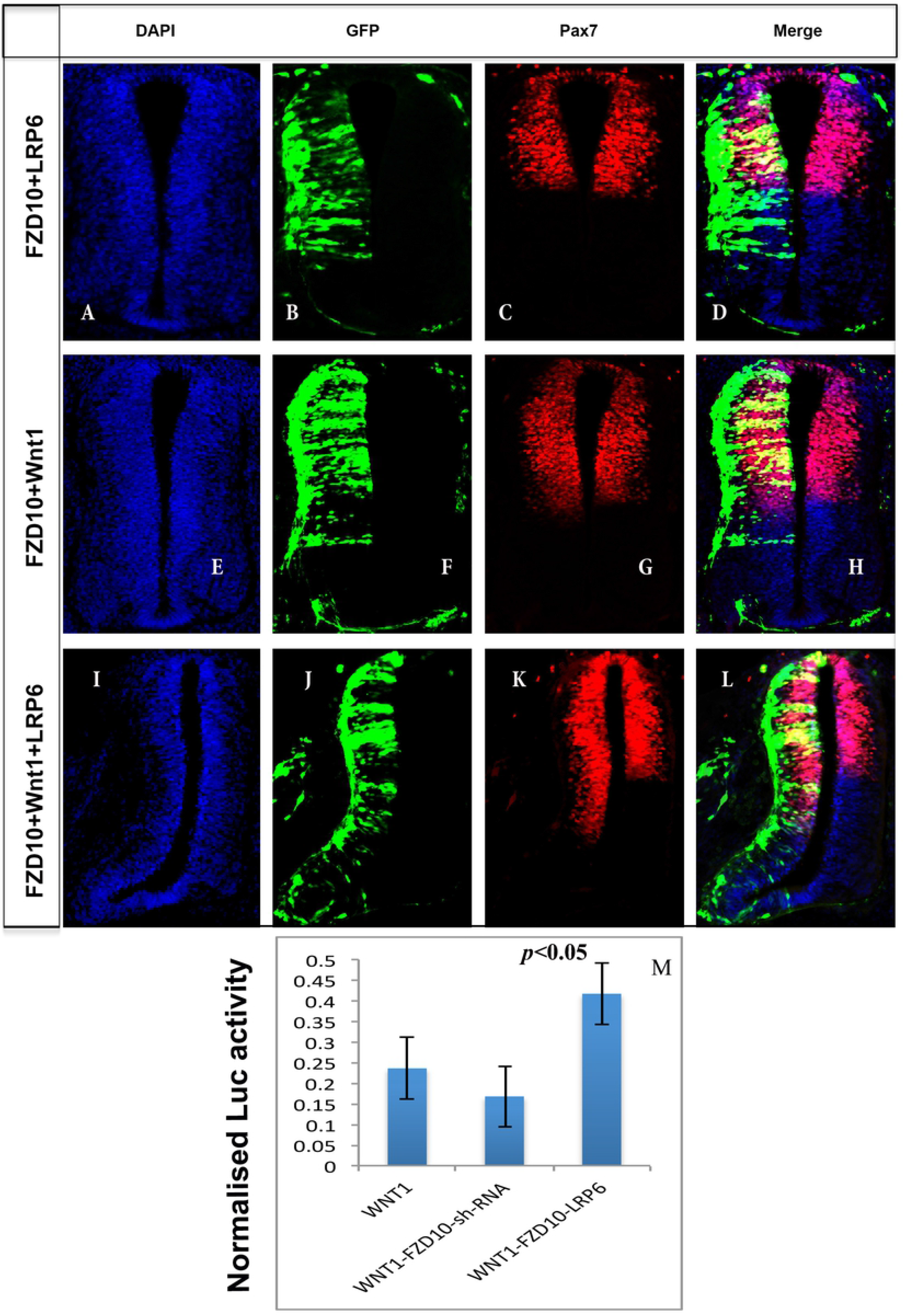
FZD10 requires the Lrp6 co-receptor to mediate Wnt1 activity in the neural tube. (A, E, I) DAPI staining of transverse sections through a HH24 neural tube. (B, F, J) GFP detection on the electroporated side reports the extend of ectopic expression of FZD10, LRP6 and Wnt1, as indicated on the left. (C, D) FZD10 co-electroporation with Lrp6 slightly restricted the ventral extend of Pax7 expression compared to the control side (n=8/10) suggesting limited availability of endogenous Wnt1 ligand. (G, H) FZD10 coelectroporation with Wnt1 attenuated the Wnt1 overexpression phenotype; the kink in the ventral spinal cord was missing and ventral Pax7 expansion was reduced (n=4/5). (I-L) Electroporation of FZD10, LRP6 and Wnt1 in combination resulted in overgrowth on the electroporated side of the spinal cord (I, J) and the ventral expansion of the Pax7 expression domain (K, L) (N=5/5). (M) Luciferase activity resulting from TOP-flash reporter expression is shown, normalized to reporter plasmid alone. The presence of Wnt1 alone increased luciferase activity, this was inhibited by FZD10 knockdown and enhanced by cotransfection of FZD10 and LRP6. All transfections also normalized to Renilla luciferase.

To quantify the Wnt activity present in the tissue we transfected a TOP-flash luciferase plasmid into the neural tube. TOP-flash luciferase reports canonical Wnt activation and was transfected either on its own or together with Wnt1, Wnt1 and FZD10 shRNA, or Wnt1, FZD10 and LRP6. Luciferase reads were normalized against Renilla and vector only reads. Wnt1 led to an increase in luciferase activity by 20%, indicative of increased transcriptional activation. This was inhibited by FZD10 shRNA, suggesting that FZD10 mediates the response to Wnt1. Addition of both FZD10 and the LRP6 co-receptor enhanced the Wnt1-induced activation of luciferase expression. These findings are consistent with the idea that Wnt1 dependent dorso-ventral patterning and neurogenesis in the developing spinal cord involves FZD10 and its co-receptor Lrp6.

## Discussion

Wnt/FZD signaling, in particular canonical Wnt signaling, is essential for neural development. This includes findings that Wnt1 and Wnt3a, two canonical Wnt ligands, are required for cell proliferation in the neural tube (Dickinson et al., 1994; Megason and McMahon, 2002; Muroyama et al., 2002). In addition, co-electroporation of Wnt1 and Wnt3a results in the ventral expansion of dorsal marker genes, Pax7 and Pax6, (Alvarez-Medina et al., 2008). Indeed, electroporation of either Wnt1 or Wnt3a leads to ventral expansion of Pax7 and Pax6 in the neural tube, indicating that both Wnt ligands can regulate dorso-ventral neural tube patterning (Supp Figs. 3 and 4). However it is not known which frizzled receptor is mediating Wnt1 and Wnt3a activity.

Here we identify FZD10 as a receptor, which mediates Wnt1 but not Wnt3a activity in the chick neural tube. We show that the dorsally restricted expression of FZD10 in the neural tube and roof plate, also reported by (Galli et al., 2014), overlaps with both Wnt1 and Wnt3a (Supp Fig. 4) and with well-characterized dorsal progenitor markers, Pax3 and Pax7, during neurogenesis (Fig. 1). In situ proximity ligation assays showed that *in vitro* FZD10 interacts with both Wnt1 and Wnt3a (Galli et al., 2014). We determined whether Wnt1 and/or Wnt3a affect FZD10 expression *in vivo*. We find that in response to Wnt1 FZD10 expression extends ventrally, but it is not affected by Wnt3a transfection into the neural tube (Fig. 4). This suggests that FZD10 expression is regulated by Wnt1 in the dorsal neural tube and is consistent with the idea that FZD10 mediates Wnt1 function in this context. Although it should be emphasized that transcriptional regulation of a FZD receptor gene by a Wnt ligand does not necessarily mean that they interact at the protein level. In the case of Wnt1 and FZD10 a direct interaction in a physiological context remains to be confirmed.

FZD10 expression suggests that it may be required for dorsal neural tube development. Consistent with this, we show that shRNA-mediated knockdown of FZD10 results in a significant decrease in the number of mitotic cells in the developing neural tube (Fig. 2). This implies that FZD10 is required for cell proliferation in the spinal cord and is in keeping with the role of canonical Wnt in cell proliferation (Dickinson et al., 1994; Megason and McMahon, 2002). Moreover, the activation of dorsal genes, Pax7 and Pax6, is inhibited by FZD10 knockdown as their expression domains are reduced (Fig. 3). Neurogenesis is also inhibited as the expression domains of differentiation markers, Lhx1/5 and Tuj1, are reduced (Fig. 3). The knockdown is mosaic, and it is not clear at present whether FZD10 affects proliferation and the expression of these marker genes directly or indirectly. In addition, FZD10 knockdown by shRNA could rescue the Wnt1-induced dorsalisation of the neural tube, as shown by lack of ventral Pax7 expansion. The effect of Wnt1 on Pax6 expansion was more subtle, but a rescue by FZD10 shRNA is still apparent (Fig. 5). In contrast, the Wnt3a mediated ventral expansion of dorsal genes, Pax6 and Pax7, is not affected by FZD10 knockdown (Fig. 5). This suggests that FZD10 is required for dorsal neural tube patterning and neurogenesis mediated by Wnt1, but not Wnt3a.

This is reminiscent of previous *in vivo* and *in vitro* studies. In particular, FZD10 did not synergise with Wnt3a in Xenopus animal cap assays, suggesting there is no interaction between Wnt3a and FZD10 (Kawakami et al., 2000). We reported previously that FZD10 synergises with Wnt1 and Wnt8, but not with Wnt3a, to induce axis duplication in Xenopus embryos (Garcia-Morales et al., 2009). In addition, FZD10-CRD did not interact with Wnt3a in co-immunoprecipitation experiments (Carmon and Loose, 2010). Thus our results agree with those of others and together they show that FZD10 mediates Wnt1 but not Wnt3a biological activity in different scenarios. In the dorsal neural tube, Wnt3a may function through different FZD receptors; good candidates are FZD1 and FZD3. Consistent with this hypothesis, Wnt3a is capable of activating canonical Wnt signalling through FZD1 in P12 cells (Chacon et al., 2008).

Interestingly, expression of FZD10 alone does not cause a ventral expansion of dorsal neural tube markers, instead the expression domains of Pax6 and Pax7 are restricted dorsally on the experimental side (Supp Fig. 5). This is surprising as FZD10 activated canonical Wnt signaling should promote proliferation and neurogenesis in the dorsal spinal cord. We propose that full-length cFZD10 may interfere in a dominant negative manner by forming ineffective receptor-ligand complexes. This is supported by a study reporting that overexpression of full-length cFZD1 or cFZD7 mimics the effect of overexpression of dominant-negative forms of these two FZDs in the developing chick wing (Hartmann and Tabin, 2000). Furthermore, excess FZD10 receptor may restrict Wnt1 ligand, for example by acting as a sponge. Consistent with this idea, FZD10 electroporation together with Wnt1 has a negative effect on the Wnt1-induced phenotype; the ventral expansion of Pax7 is less pronounced compared to that seen after Wnt1 electroporation alone (Fig. 6G, compare with Fig. 5A). Our data suggest that FZD10 requires Lrp6 co-receptor to activate canonical Wnt signaling effectively. Lrp6 is expressed in the developing chick neural tube and binds Wnt1 (Avile and Stoeckli., 2015; He et al., 2004; MacDonald and He et al., 2012; Tamai et al., 2000). Co-transfection of Lrp6 with FZD10 and Wnt1 into the neural tube leads to a dramatic lengthening of the dorso-ventral axis and to ventral expansion of the Pax7 expression domain compared to the control side (Fig. 6 I-L). The overgrowth on the experimental side is likely due to increased proliferation, leading to increased cell number. In addition, we show that Wnt1 increased the luciferase activity of a canonical Wnt reporter by 20%, indicative of increased transcriptional activation. Addition of both FZD10 and the LRP6 co-receptor further enhanced the Wnt1 induced activation of luciferase expression, but FZD10 shRNA negatively affected the response. (Fig. 6M).

Taken together, we show that FZD10 is required for neural tube development and we propose that it may be the cognate receptor that mediates Wnt1 biological activity in the developing neural tube, although direct interactions remain to be confirmed *in vivo*. In addition, we show that Lrp6 is essential for effective signaling to regulate proliferation and dorso-ventral patterning. At present Wnt/FZD interactions and ligand-receptor selectivity are not fully understood. We propose that the chick neural tube presents an accessible tool to dissect Wnt-FZD interactions and selectivity in more detail.

## Materials and Methods

### Injection and electroporation into neural tube

Fertile chick eggs were obtained from a commercial supplier and incubated for up to 3 days. Embryos at this stage are not subject to any legislation (Animals Scientific Procedures Act) as they are less than 2/3 of gestation. The research conducted in the laboratory was approved by the UEA Animal Welfare & Ethical Review Board (08032018). Fertilized eggs were incubated at 37°C until the desired stage of development was reached (Hamburger and Hamilton, 1992). Expression constructs or shRNA were injected into the lumen of neural tubes of HH11-12 embryos and embryos were electroporated using 24V, five 50msec pulses with 100msec intervals. Embryos were harvested after 24 or 48 hours for analysis, at least 3 embryos were examined per experimental condition and marker gene.

### Whole-mount in situ hybridization, cryosections, area measurements and photography

Embryos were collected into DEPC treated PBS, cleaned and fixed in 4% paraformaldehyde overnight at 4°C. Whole mount in situ hybridization was performed as previously described (Schmidt et al., 2004; Goljanek-Whysall et al., 2014). For cryosectioning, embryos were embedded in OCT (Tissuetec) and 20 □m sections were collected on TESPA coated slides, washed with PTW, coverslipped with Entellan (Merck, Germany) and examined using an Axioplan microscope (Zeiss). Whole mount embryos were photographed on a Zeiss SV11 dissecting microscope with a Micropublisher 3.5 camera and acquisition software. Sections were photographed on an Axiovert (Zeiss) using Axiovision software. Images were imported into Fiji/ImageJ, and areas of staining were calculated from binary images by calculating pixel numbers from injected and noninjected sides (Abou-Elhamd et al., 2015; Mok et al., 2018). A minimum of 10 sections from three embryos were analysed for each experiment. Statistical analysis used GraphPad Prism (version 6) software. Mann–Whitney nonparametric two-tail testing was applied to determine P-values. Montages of images were created and labeled using Adobe Photoshop.

### DNA constructs

Plasmids encoding mouse Wnt1 and Wnt3a (pCIG) were kindly provided by Elisa Marti (Alvarez-Medina et al., 2008). The full length coding sequence for FZD10 was amplified by PCR from cDNA prepared from HH18 chick embryos using standard molecular biology protocols. Primers were designed using FZD10 sequences for the chicken from NCBI (http://www.ncbi.nlm.nih.gov) using accession number (NM_204098.2). Restriction sites were added, FZD10 primer sequences were: Not1+FZD10 forward: 5’-GCGGCCGCATGTGCGAGTGGAAGAGGTG-3’ and EcoR1+ HA tag +FZD10 reverse:5’-GAATTCTCAAGCGTAATCTGGAACATCGTATGGGTATCATACACAG-GTGGGTGGTTG-3’. PCR products were cloned into pGEM-T (Promega) and sequenced. FZD10 was subcloned into the pCA-IRES-GFP vector using EcoRI and NotI restriction enzymes for electroporation. pCS2-hLrp6 was obtained from Addgene. Short RNA hairpin (sh-RNA)-based expression vectors for RNA interference pRFP-C-RS (FZD10 shRNAs and scrambled shRNA) were purchased from Origene. The three sequences were: ‘A’–GTACAACATGACGAGAATGCC-GAACCTGA,‘B’-TGGATTGCCATCTGGTCCATTCTGTGCTT, ‘C’-GCAAGCGTTATTACCAGT-AGT GGAATCTA

### In vivo luciferase-reporter assay

Transcriptional activity assays of β-catenin/Tcf pathways were performed in the neural tube as described by (Alvarez-Medina et al., 2008). Chick embryos were electroporated at HH stage 11/12 with the following DNAs: Wnt1, FZD10, hLrp6 and FZD10 shRNA, or with empty pCA vector as control, together with a TOPFLASH luciferase reporter construct containing synthetic Tcf-binding sites (Korinek et al., 1998) and a Renilla-luciferase reporter (Promega) for normalization. Embryos were harvested after 24 hours incubation and GFP-positive neural tubes were dissected and homogenized with a douncer in Passive Lysis Buffer. Firefly- and Renilla-luciferase activities were measured by the Dual Luciferase Reporter Assay System (Promega). Statistical analysis was performed by Student’s t-test.

### Immunohistochemistry

Immunohistochemistry was performed as described previously (Abu-Elmagd et al., 2010). Sections were incubated overnight at 4°C with primary antibodies at the following concentrations: Pax6, Pax7, Nkx2.2 (74.5A5), Islet1 (40.2D2), Lhx1/5 (4F2) (1:100, all from Developmental Studies Hybridoma Bank, University of Iowa), anti-rabbit phospho-histone H3 (5:1000, Abcam), anti-mouse Tuj1 (2:1000, Covance), anti-rabbit RFP (2:1000, Abcam). Secondary antibodies were anti-rabbit Alexa Fluor 488/568, anti-mouse Alexa Fluor 488/568, and anti-mouse Alexa Fluor 350 (Invitrogen) at 1 mg/ml in 10% goat serum/PBS. DAPI was used at a concentration of 0.1 mg/ml in PBS. After staining, cryosections were mounted and visualized using an Axioscope microscope using Axiovision software (Zeiss, Germany). Images were imported into Adobe Photoshop for analysis and labeling. Statistical analysis was performed by Student’s t-test.

## Acknowledgements

We thank Drs Timothy Grocott and Gi Fay Mok and the rest of the Wheeler and Münsterberg labs for discussions, Prof Elisa Marti for providing plasmids and insightful comments, Dr Paul Thomas for support in the Henry Wellcome Laboratory of Cell Imaging.

## Author contributions

Conceptualization: AM, GW; Methodology: AFA, AM, GW; Validation: AFA, Formal analysis: AFA, AM, GW; Investigation: AFA, AM, GW; Resources: AM, GW; Data curation: AFA, AM, GW; Writing: AFA, AM, GW; Visualization: AFA, AM, GW; Supervision: AM, GW; Project administration: AM, GW; Funding acquisition: AFA, AM, GW

